# *c-fos* induction in the choroid plexus, tanycytes and pars tuberalis is an early indicator of spontaneous arousal from torpor in a deep hibernator

**DOI:** 10.1101/2023.12.19.572338

**Authors:** Fredrik A.F Markussen, Fernando Cázarez-Marquez, Vebjørn J. Melum, David Hazlerigg, Shona Wood

**Affiliations:** Arctic seasonal timekeeping initiative (ASTI), Arctic Chronobiology & Physiology, Arctic & Marine Biology, BFE, UiT - Arctic University of Norway, Tromsø, Norway

**Keywords:** Hibernation, Thermoregulation, *c-fos*, Golden hamster, choroid plexus, tanycytes, dorsomedial hypothalamus, pars tuberalis

## Abstract

Hibernation is an extreme state of seasonal energy conservation, reducing metabolic rate to as little as 1% of the active state. During the hibernation season, many species of hibernating mammals cycle repeatedly between the active (aroused) and hibernating (torpid) states (T-A cycling), using brown adipose tissue (BAT) to drive cyclical rewarming. The regulatory mechanisms controlling this process remain undefined but are presumed to involve thermoregulatory centres in the hypothalamus. Here, we use the golden hamster (*Mesocricetus auratus*), and high-resolution monitoring of BAT, core body temperature (T_b_), and ventilation rate, to sample at precisely defined phases of the T-A cycle. Using *c-fos* as a marker of cellular activity we show that although the dorso-medial hypothalamus (DMH) is active during torpor entry, neither it nor the pre-optic area (POA) show any significant changes during the earliest stages of spontaneous arousal. Contrastingly, in 3 non-neuronal sites previously linked to control of metabolic physiology over seasonal and daily timescales, the choroid plexus (CP), pars tuberalis (PT) and third ventricle tanycytes, peak *c-fos* expression is seen at arousal initiation. We suggest that through their sensitivity to factors in the blood or cerebrospinal fluid (CSF), these sites may mediate metabolic feedback-based initiation of the spontaneous arousal process.

**Summary statement:** Non-neuronal sites linked to control of daily and seasonal metabolic physiology, peak in *c-fos* expression during arousal from torpor, suggesting metabolic feedback-based initiation of the spontaneous arousal process.

## Introduction

Hibernation is a seasonally regulated process of energy conservation achieved through extreme shut down of metabolic energy expenditure, sometimes to as little as 1% of that in the euthermic state, allowing species to survive extended winter periods of low energy availability (Kenagy et al., 1989; Mckee and Andrews, 1992; Ruf and Geiser, 2015). Hibernation is characterised by multi-day bouts of torpor during which metabolic rate is suppressed and core body temperature (T_b_) falls, in some species to within a few degrees of ambient temperature (T_a_), while breathing and heart rate slow to an extent not expected to support life in endothermic organisms (Lyman et al., 1982).

In many hibernating species, torpor bouts are separated by relatively short arousal episodes during which T_b_ rises back to euthermic levels, so that the hibernation season as a whole comprises a series of torpor – arousal (T-A) cycles. This presents a paradox, why would an organism undergo a costly arousal if the ultimate goal of hibernation is to save energy? Since homeostatic processes consume energy to maintain healthy cellular environments, one scenario is that arousal episodes allow restoration of healthy cellular conditions by reactivating aspects of homeostasis that are compromised or suspended during torpor.

While this narrative provides plausible ultimate reasoning for the evolution of periodic arousal, it does not address the proximate mechanisms by which T-A cycling is controlled. Here, two basic models can be considered: a clock-based mechanism and a metabolic feedback-based mechanism. In its most explicit formulation, the former is conceived as an extension of circadian function (Körtner and Geiser, 2000; Malan, 2010). The hypothalamic circadian pacemaker in the suprachiasmatic nucleus (SCN) controls daily T_b_ cycles in euthermic mammals and the expression of daily torpor episodes in Siberian hamsters (Körtner and Geiser, 2000; Ruby and Zucker, 1992; Ruf, T., Steinlechner, S., and Heldmaier, 1989). A role for circadian control mechanisms in multiday T-A cycling is not favoured, however. Except for the first torpor bout of the season, T-A cycles show no circadian organisation (Hut et al., 2002; Williams et al., 2012; Williams et al., 2017), and SCN molecular clock gene oscillations flatten out (Ikeno et al., 2017; Revel et al., 2007). Malan proposed a ‘poorly temperature compensated circadian mechanism’ in which slow running of the circadian oscillation at low temperatures determines timing of arousals (Malan, 2010; Malan et al., 2018), but this conjecture fails to account for changes in T-A cycle periodicity, particularly at the beginning and end of the hibernation season, or for the finding that the period of T-A cycling is the same in wild-type Syrian hamsters and in *tau* mutant hamsters in which the circadian clock runs fast (Oklejewicz et al., 2001). Hence current models focus on the second possibility, namely that T-A cycle regulation stems from accumulation (or depletion) during the torpid state of metabolite(s) which act as signals to stimulate the arousal process, and that levels of these metabolic signals are then reset during arousal permitting torpor re-entry (reviewed in:(van Breukelen and Martin, 2015)). Since arousal is initiated by combined increases in pulmonary-cardio-vascular activity and brown adipose tissue (BAT) activation, this framing places an emphasis on metabolic signals that access brain mechanisms controlling sympathetic stimulation of these tissues.

A study in the Siberian chipmunk identified a “hibernation factor”, produced in the liver and transported to the brain via the choroid plexus (Kondo et al., 2006), indicating a link between peripheral communication to the brain via the circulation and cerebrospinal fluid (CSF) in the regulation of hibernation. However, this study did not investigate changes in hibernation factor levels during T-A cycling, instead emphasizing a role as a putative seasonally permissive factor (Kondo et al., 2006). The circuits controlling BAT thermogenic activity in euthermic rodents include the temperature-sensitive pre-optic area (POA), which acts via the dorsomedial hypothalamus (DMH)(Cannon and Nedergaard, 2004; Morrison, 2016). In mice, chemo/opto-genetic manipulation of the activity of neurons within the POA can suppress T_b_ and metabolic rate (Hrvatin et al., 2020; Takahashi et al., 2020; Upton et al., 2021) to produce a torpor-like state, suggesting that metabolic feedback to these sites could be involved in natural torpor regulation.

To identify key signals controlling spontaneous T-A cycling, it is necessary to overcome the challenge of repeatable sampling immediately prior to, and in the earliest stages of spontaneous arousal from torpor. Because this is difficult, “forced” arousal by handling or changing room temperature has been employed (Regan et al., 2019). Unfortunately this necessarily alters the dynamics of the T-A cycle and causes stress (Tähti and Antti, 1977; Utz and van Breukelen, 2013), potentially confounding attempts to characterize brain mechanisms involved in spontaneous T-A cycling. Bratincsak *et al*. combined T_b_ measurement and expression of the immediate early gene *c-fos* (Bratincsák et al., 2007) to explore brain changes during spontaneous T-A cycles in thirteen lined ground squirrels. While this revealed *c-fos* changes in many brain sites, the use of T_b_ as the sole indicator of arousal status inevitably misses the early stages of the arousal process, which may precede core T_b_ by almost one hour (Markussen et al., 2020).

Here, we have developed an enhanced telemetric approach based on BAT temperature combined with T_b_ and ventilation rate to characterize T-A cycling at high temporal and temperature resolution in hibernating golden hamsters, a species showing highly reproducible T-A cycles (Chayama et al., 2016; Lyman, 1948; Lyman et al., 1982). Using *c-fos* RNA expression, we sought to capture the early dynamics of brain activation during spontaneous T-A cycling. Our data reveal changes in the choroid plexus, pars tuberalis and tanycytes lining the 3^rd^ ventricle of the hypothalamus, as the earliest brain indicators of initiation of spontaneous arousal.

## Materials and Methods

### Animals & Experimental set up

All experimental procedures were carried out at the animal facility of Arctic Chronobiology and Physiology at UiT – the Arctic University of Norway. All procedures and experiments were approved under national legislation by the Norwegian Food Safety Authority under the laboratory animal administration’s supervision and application system ID: 24904. For 4 weeks, 5 months old male golden hamsters (n=30, *Mesocricetus auratus,* RJHan:AURA) purchased from Janvier Labs (Le Genest-Saint-Isle, France) were kept in sibling groups with *ad libitum* access to food and water, housed under long photoperiod (LP: 16L:8D) and 21°C. Prior to induction of hibernation, Anipill® temperature loggers (Animals-monitoring, Hérouville Saint-Clair, France) and IPTT-300 (BMDS/Plexx BV,Netherlands) tags were implanted under surgical anesthesia in the abdominal cavity and interscapular brown fat deposit, respectively. The animals were then single housed and allowed 2 weeks to recover in LP. To induce hibernation, single housed hamsters were switched to short photoperiod (SP: 8L:16D) for 4 weeks at 21°C (SP-warm) before lowering the ambient temperature to 6.5 °C (SP-cold).

### Physiological and behavioural monitoring

To reduce stress and disturbance to the animals during the SP-cold phase, a select group of trained personnel conducted daily monitoring, thereby maintaining a stable and controlled environment conducive to animal welfare.

Real-time monitoring of interscapular BAT temperature (T_iBAT_) was done using a mix of proprietary hardware and readily available electronics components using open-source practices, together with an open-source software stack. Iterative reads of T_iBAT_ were done using BMDS DAS-8027 (BMDS/Plexx BV, Netherlands) reader, and fixed in position near the scapular region of the hibernating hamster. The reader was configured to operate in automatic scanning mode, recording the IPTT-300 every 3-5 seconds. Data acquisition from the reader was wirelessly transmitted through BMDS communications module to a computer in a separate room. The data was written into a text document using Atom text editor (https://github.com/atom/atom) and automatically saved on change. A program written using NodeRED v.19.9.0 (OpenJS Foundation) monitors this file and automatically extracts the last row of information as soon as data is written, formats it, and pushes the data into an InfluxDB v.1.8.1 (InfluxData Inc, San Francisco, CA, USA) database housed in the local computer (JavaScript for replication of the NodeRED program is available here: https://github.com/ShonaWood/cFOSGHam). Ambient temperature (T_a_) was also continuously monitored using RuuviTag Bluetooth sensors (Ruuvi Innovations Ltd, Riihimäki, Finland), data were pushed to Influx DB using NodeRed. InfluxDB data was accessed and displayed using Grafana v.10.0.1 (GrafanaLabs, New York, NY, USA), this allowed real-time monitoring from any computer or smart phone. We established a torpor baseline T_iBAT_ temperature by averaging T_iBAT_ values below 10`°`C and subtracting the T_a_ to calculate the thermal distance (TD). Subsequently we calculated the change from this baseline (ΔT), therefore as the animal remains torpid the value remains at zero, even with ambient temperature fluctuations. Mathematically this is simply expressed as:

ΔT = T_iBAT_ - (T_a_ + TD) | TD = mean(T_iBAT_ < 10) - T_a_).

Departures from ΔT were then used to detect animals spontaneously arousing.

In parallel an infrared sensitive camera (Raspberry Pi Camera Module 2 NoIR, Raspberry Pi Foundation, UK) connected via Raspberry Pi (model 3B+, Raspberry Pi Foundation, UK) was used to stream a video feed to monitor and record animal activity and respiration rate using software developed by (Singh et al., 2019).

We then analyzed twelve T-A cycles from 6 individuals to select physiological sampling criteria with the goal of reducing inter-individual variation and identify the earliest possible reliable predictors of arousal. Our criteria were; Animals entering torpor (ENT<25`°`C) must demonstrate a minimum of a 12-hour inter bout euthermic (IBE) duration, followed by a steady decrease in observed anipill T_b_ to below 25`°`C. Additionally the IPTT read must be below 25`°`C and the animal must be in curled up torpid posture. Torpid animals (T-40h) were sampled 40 hours after ENT<25`°`C, if Tb and T_iBAT_ were within 2°C of T_a,_ and ventilation frequency (VF) was less than 2 per minute. Arousal 1 (AΔ0.5`°`C) was defined only if an animal had entered torpor and remained torpid for longer than 40 hours, and subsequently showed a VF >10 per minute and a 0.5`°`C change in T_iBAT_ from torpor baseline (ΔT). Arousal 2 (AΔ3`°`C) followed the same criteria as arousal 1 but was sampled at ΔT=3`°`C. IBE was defined as an animal that had entered torpor, remained in torpor for longer than 40 hours, subsequently aroused and then reached a T_b_ of 30°C. Animals were sampled 1 hour (IBE-1h) and 12 hours (IBE-12h) after euthermia was reached.

Prior to sampling all animals had shown at least 3 full T-A cycles, these data were used to verify the sampling criteria prior to sampling in the 4^th^ T-A cycle. On reaching the above criteria the animals were euthanized by cervical dislocation. The brain was rapidly dissected and frozen in isopentane before storage at −80°C until sectioning.

### *In-Situ* hybridization

Regions of interest, preoptic area (POA, Bregma ca. 0.8 to 0.95) and mediobasal hypothalamus (MBH, Bregma ca. −2.5 to −2.8), were collected in 14 µm sections onto Fisherbrand Superfrost Plus slides using a Leica CM3050S cryostat and stored at −80°C until further processing.

On the day of hybridization, sections were dried for 15 min at 50°C before fixation with 4% formaldehyde solution for 20 min. Sections were acetylated and delipidated using 1% (volume/volume (v/v)) triethylamine (Sigma, 90335) with 0.25% (v/v) acetic anhydride (Sigma, 539996) and 0.1% (v/v) Triton-X-100 (Sigma, X100) in nuclease-free water, respectively. The sections were hybridized with a 200 ng/mL *c-fos* antisense probe (A gift from Valerie Simonneaux and Paul Klosen, University of Strasbourg, 793-1193 of Genbank accession: XM_005086369.4) overnight at 58°C in hybridization buffer containing 5x Denhardt’s solution (Sigma, D2532), 0.25 mg/mL yeast tRNA (Roche, 10109525001), 0.2 mg/mL fish sperm ssDNA (Roche, 11467140001), 5x saline-sodium citrate (SSC, Sigma, 8310-OP), and 50% (v/v) formamide (Sigma F9037) in nuclease-free water (Sigma, F9037). Following hybridization, washes were made in decreasing concentrations of SSC (5x and 2x) at 55°C, 50% formamide in 0.2x SSC at 55°C, and 0.2x SSC at room temperature. After blocking using 1% Roche blocking solution (Roche, 11096176001), sections were incubated for 3 hours with Anti-Digoxigenin-AP Fab fragments (dilution 1:3000) (Roche, 11093274910) at room temperature, followed by washes in A-dig and alkaline phosphate buffer. The signal was then visualized by incubating the slides for 48 hours in a solution containing 0.33 mg/mL NBT (Nitro-blue tetrazolium chloride, Roche, 11383213001) and 0.165 mg/mL BCIP (5-Bromo-4-chloro-3-indolyl phosphate, Roche, 11383221001) in alkaline phosphatase buffer. Once the signal had developed properly, the slides were prepared for immunohistochemical processing to visualize vimentin and cell nuclei. After three five-minute washes in PBS pH 7.6 containing 0.05% (v/v) Tween-20 (PBS-Tw, Sigma, P1379), the slides were incubated in blocking buffer containing PBS-Tw, 0.1% (weight/v) cold water fish skin gelatin (Sigma, G7041), 0.1% (w/v) bovine serum albumin (Sigma, A2153), and 3 mM glycine (Sigma, G7126) for one hour at room temperature, followed by overnight incubation at 4°C in blocking buffer containing mouse anti-vimentin antibody (Sigma, MAB3400) diluted 1:2000. After four 10-minute washes in PBS-Tw at room temperature, the secondary antibody was applied by incubating the slides for 1 hour at room temperature with blocking buffer containing Alexa Fluor 647-conjugated goat anti-mouse antibody (1:1000, Sigma, SAB4600354). The slides were then briefly rinsed in PBS-Tw before a 10-minute incubation in PBS-Tw containing 1:2000 SYTOX Orange (Thermo Fisher, S11368), followed by four 10-minute washes in PBS-Tw and a final rinse in PBS. The slides were then cover slipped and sealed using nail polish and 1.5H coverslips over antifade solution containing 90% (v/v) glycerol (Sigma, G7757), 10% (v/v) PBS, and 2.5% (w/v) diazabicyclo 2.2.2 octane (DABCO, Sigma, 290734)

The slides were scanned using an Olympus VS120 slide scanning microscope at the core microscopy facility at UiT. Overview images of the sections were obtained using a Cy3 filter set, then regions of interest were marked up in the OlyVIA software and, by using the x40 objective, high-resolution images were obtained. The images were analyzed using QuPath v0.4.3 by annotating regions of interest (ROIs). The number of cells within each ROI was detected and quantified using the SytoxO channel in the standard cell detection plugin in QuPath. To quantify the amount of *c-fos* within each ROI, a threshold was used to select the signal, and the *c-fos*-positive area within the ROI was quantified by threshold. Cells are then classified as either positive or negative depending on the intensity above threshold within cell boundaries. The percentage of *c-fos* positive cells within each ROI was calculated. Figures were made by exporting ROIs to ImageJ v1-54f where they were assembled using FigureJ package.

#### Statistical analysis

The physiological and cell counts data were analyzed in R version 4.3.1. The scripts for the analysis and the data are available on github: https://github.com/ShonaWood/cFOSGHam

Time of day preference was assessed using Rayleigh test of circular uniformity using R. Significant differences between numbers of *c-fos* positive cells was assessed by one-way ANOVA and post-hoc testing by Dunnett’s multiple comparisons test using R.

## Results

### Reproducible induction of hibernation and reliable prediction of torpor entry and arousal

We switched animals from a long photoperiod (16L:8D) at 21°C (LP) to a short photoperiod (8L:16D) at 21°C (SP-warm) for 4 weeks, then we lowered the temperature to 6.5°C (SP-cold) and monitored core body temperature (T_b_) continuously (Figure 1A). Within 8 weeks of exposure to SP-cold conditions, 80% of our animals had commenced multi-day T-A cycling, this increased to 93% by 12 weeks (Supplementary Figure 1A). Similar to previous observations, we observed a gradual resetting of mean T_b_ from 36.4°C (Standard deviation (SD):0.83°C) to 33.4°C (SD:0.98°C) after 4 weeks in SP-cold, indicating a seasonal preparatory phase prior to the expression of hibernation (Chayama et al., 2016)(Figure 1A & Supplementary Figure 1B). We also observed so-called ‘test-dropś (Figure 1A, B)(Sheriff et al., 2012) in 86% of our animals prior to the initiation of T-A cycling. During test drops the mean T_b_ is 21.4`°`C (SD: 7.35) and the mean test-drop duration was only 5.3 hours. Both entry to, and, arousal from test drops showed circadian organisation (Supplementary Figure 1C). Not all animals showed test-drops prior to multiday T-A cycling, indicating that they are not a pre-requisite for hibernation.

**Figure 1:**
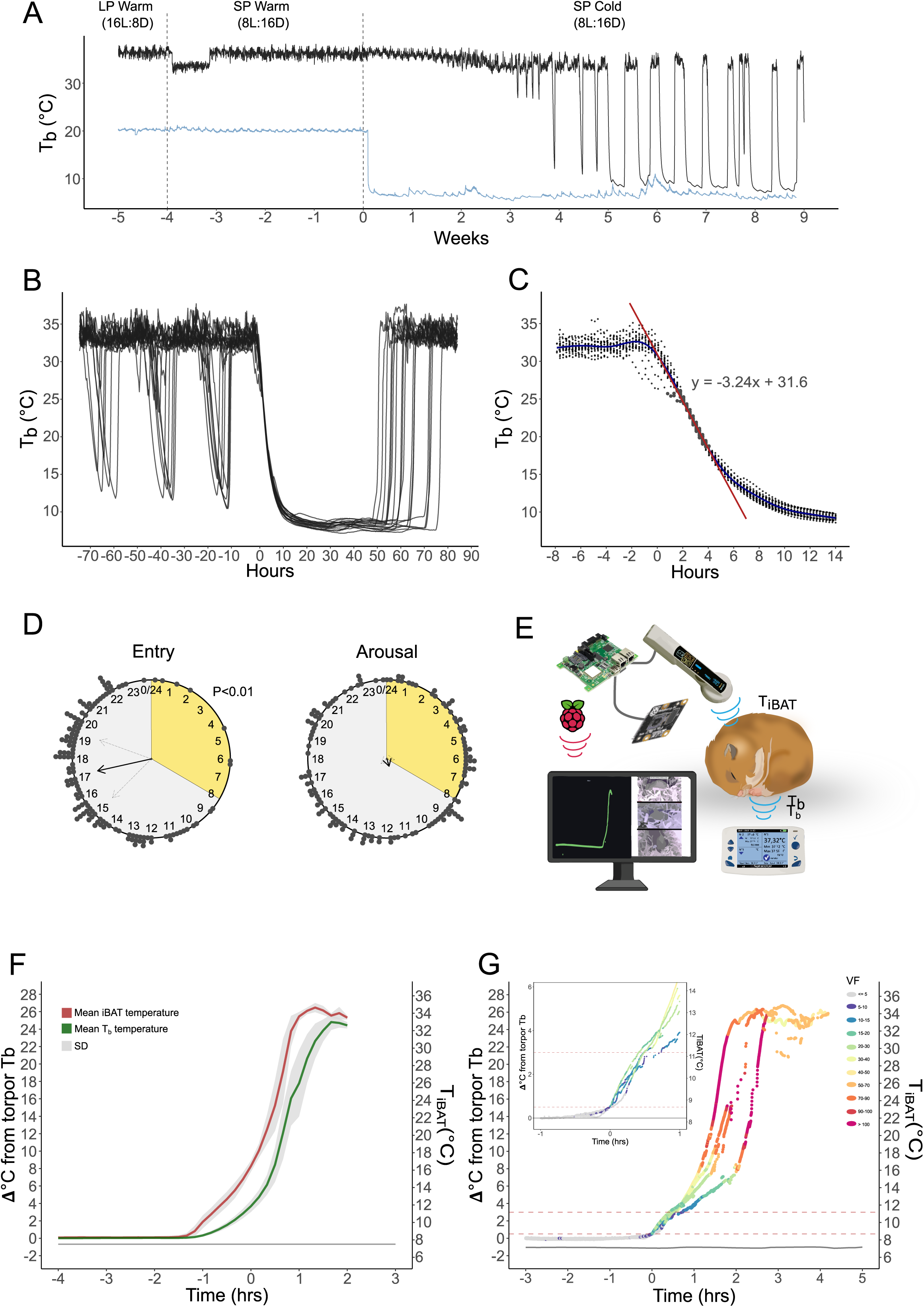
Physiological monitoring of the golden hamster during hibernation. A. Representative core body temperature trace (T_b_ `°`C, black line) and ambient temperature (T_a_ `°`C, blue line) over the whole experiment. Hamsters were kept on long photoperiod (LP (16L:8D), T_a_ = 21°C) for 6 weeks then, moved to short photoperiod and warm conditions (SP= 8L:16D, T_a_ = 21°C) for 4 weeks. Finally, the T_a_ was lowered to 6.5°C for the remainder of the experiment. The vertical dotted lines indicate when the changes in light or temperature conditions occurred. Note the drop in Tb in the first week of SP, this was due to the anipill shifting into the testis sack, which resolved itself. B. Body temperature (T_b_`°`C) traces of 18 hamsters synchronized to first torpor bout entry, illustrating test-drops prior to T-A cycling and first T-A cycle. C. Body temperature (T_b_ `°`C) of entry to torpor from 18 individuals to explore variation in entry dynamics. Red line is a regression analysis for the steepest part of the slope, showing an average maximal cooling rate of −3.24°C per hour. D. Raleigh plots showing the time of entry and arousal from 28 individuals. Photoperiod is represented by grey for dark and yellow for light. The black dots are an individual arousal or entry event. The black arrow indicates the mean time of entry or arousal, and the light grey arrows indicates the standard deviation. The length of the arrow represents mean resultant length as an indicator of concentration around the mean, therefore reflecting the statistical significance. The Raleigh statistical test was used to test for a time-of-day preference in arousal and entry. Only the entry was significant p<0.01. E. A schematic of the remote monitoring system for arousal from torpor. IPTT-300 was used to monitor iBAT temperature dynamics. A live video feed using a raspberry pi and camera module measured ventilation frequency (VF). iBAT temperature is continuously read every 3-5 seconds and all data is parsed to a networked database immediately accessible to local networked devices. The computer screen in the schematic shows the video feeds for VF and the green line is the iBAT temperature from an animal that has completed an arousal. In addition, Anipills were used to monitor core body temperature. F. The change in iBAT temperature (red line) and core body temperature (green line) from torpor baseline from 10 representative arousals (Left axis Δ`°`C). Right axis shows actual iBAT temperature. Mean values are shown with standard deviations in grey shading. G. A closer analysis of iBAT temperature dynamics during arousal of four representative arousals. The colour of the dots reflects ventilation rate. The change in iBAT temperature from torpor baseline is plotted on the left axis (Δ`°`C) and the actual temperature of the iBAT is plotted on the right axis (`°`C). The dashed red line indicates 0.5°C and 3°C change from torpor baseline. Inset graph top left: Zoom in on the first hour of arousal.

Once initiated, T-A cycles were reproducible within and between animals, with T_b_ falling to within an average of 0.75°C of T_a_. Torpor bout length increased from 56.5 hours (SD: 9.9) at the first bout to a maximum of 75.5 hours (SD: 8.9) by the sixth bout, while the interbout euthermic (IBE) interval decreased (from 61.2 hrs (SD:38.3) to 22.9 hrs (SD:7.86), Supplementary Figure 1D).

The T_b_ dynamics of the entry phase to torpor follows an inverse sigmoidal trajectory (Figure 1C). This is somewhat variable in between 33 – 26°C, with some animals showing a “step-wise” entry pattern; thereafter Tb declines approximately linearly at a max cooling rate of −3.24`°`C per hour, until about 15°C when the approach to nadir values slows (Figure 1C). Initiation of torpor is more likely to occur in the dark phase (approx.17 to 22 hours after lights on (p<0.01 by Rayleigh test of circular uniformity, Figure 1D).

Consistent with previous studies, arousal from torpor showed no circadian organisation (Figure 1D) (Hut et al., 2002; Oklejewicz et al., 2001; Williams et al., 2012; Williams et al., 2017). Since T_b_ is not a good early indicator of arousal (Markussen et al., 2020), we used IPTT-300 tags to measure interscapular BAT temperature (T_iBAT_) every 3-5 seconds, and monitored ventilation frequency (VF) by video-streaming (Figure 1E). During spontaneous arousal, increases in T_iBAT_ precede T_b_ by up to 40 minutes (Figure 1F). We noted some variability in the arousal process when T_iBAT_ was between 4`°`C and 10°C from the torpor baseline, after which ventilation rates rapidly increased towards peak values together with the max rewarming speed of T_iBAT_ and T_b_ (+18.3 °C / h). Despite variability in its initiation, arousal was completed within 2.5-3 h in all animals (Figure 1G). To determine the earliest possible time at which we could reliably predict an arousal event and clearly differentiate it from animals thermoregulating during torpor, we monitored 12 separate arousals. We found that an increase in T_iBAT_ of 0.5°C above torpor baseline coupled with a VF > 10 was the earliest reliable predictor of a forthcoming arousal event, approximately 40 minutes before a detectable change in T_b_ (Figure 1F, G).

### The DMH expresses *c-fos* during entry to torpor

We used RNA expression of the immediate early gene *c-fos* by in-situ hybridization and validated our probes using a light pulse paradigm and staining of the suprachiasmatic nucleus (Supplementary Figure 2A)(Earnest et al., 1990). Then we sampled animals that had undergone at least 3 full T-A cycles at the following phases (for details of calculation see Methods): Entering torpor (ENT<25`°`C), 40 hours torpid (T-40h), the earliest spontaneous arousal (AΔ0.5`°`C), early spontaneous arousal prior to phase of high inter-animal variation (AΔ3`°`C), and interbout euthermic animals after 1 hour (IBE-1h) and 12 hours (IBE-12h) (Figure 2A).

**Figure 2:**
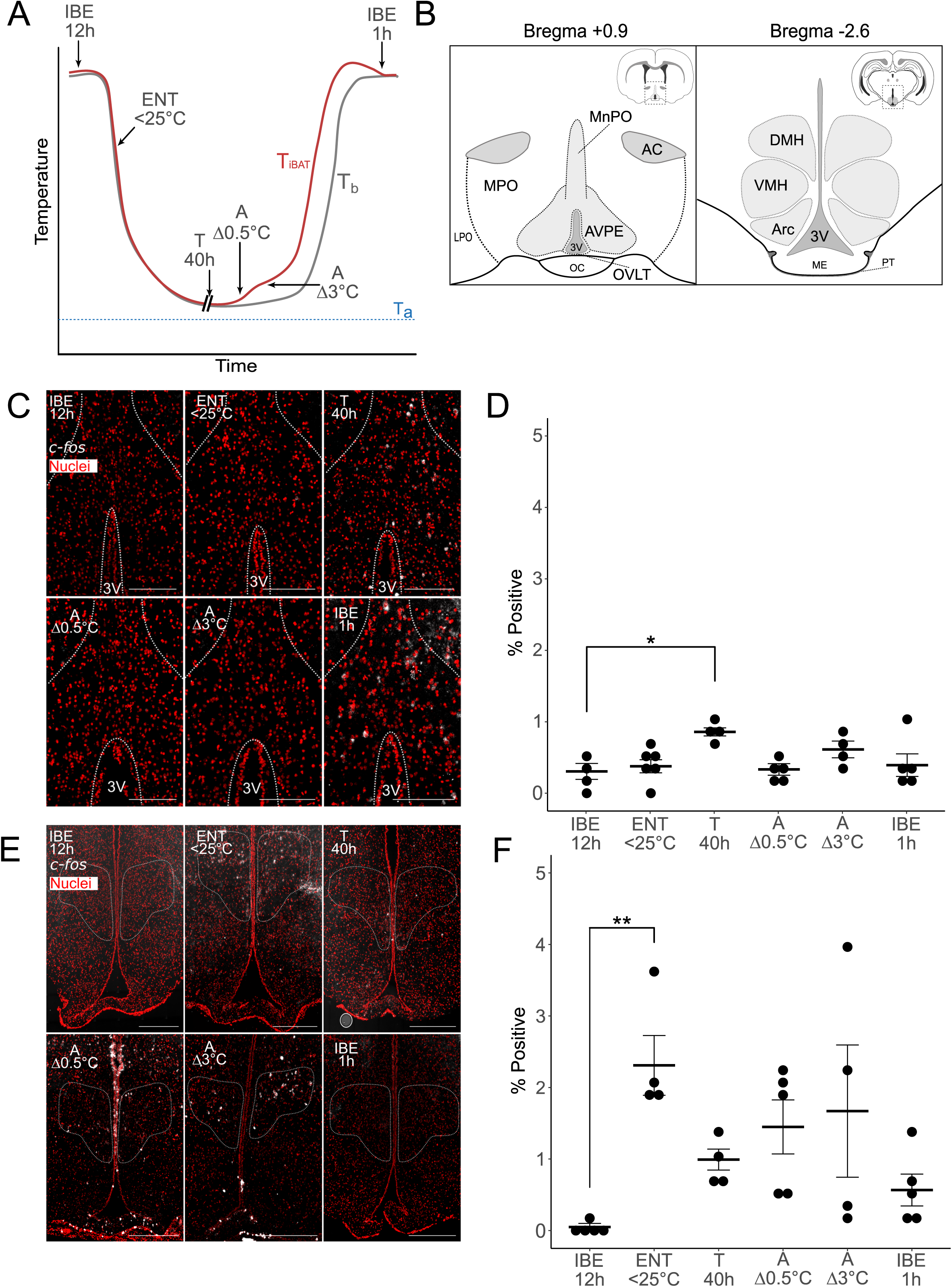
*c-fos* expression in the POA and DMH during T-A cycling. A. A schematic showing interscapular BAT temperature (T_iBAT_, red) and core body temperature (T_b_, grey) and the sampling points taken during the T-A-cycle. Interbout euthermic animals sampled either 1 or 12 hours after arousal (IBE-1h, IBE-12h). Entry to torpor animals sampled when T_b_ and T_iBAT_ are less than 25°C (ENT<25°C). Torpid animals sampled 40 hours into the torpor bout (T-40h). Early arousal animals defined by deviation in T_iBAT_ from the torpor baseline of either 0.5°C or 3°C (AΔ0.5°C and AΔ3°C). B. Brain regions of interest defined by the golden hamster brain atlas. Left panel: Bregma +0.9 in the pre-optic hypothalamic area (POA): MnPO; median preoptic area, AVPE; anteroventral periventricular area, MPO; medial preoptic area, LPO; lateral preoptic area, OVLT; organum vasculosum lamina terminalis, OC; optic chiasm, AC; anterior commissure. Right: Bregma - 2.6 in the medio basal hypothalamus: DMH; dorsomedial hypothalamus, VMH; ventromedial hypothalamus, Arc; Arcuate nucleus, 3V; third ventricle, ME; median eminence, PT; Pars tuberalis. C. Representative images of in-situ hybridization using *c-fos* (white) and sytox orange to show nuclei (red) in the preoptic area (POA), indicated for white outline, for each group defined in A. Scalebar 500 µm. D. Quantification of the percentage of positive cells in the preoptic area (POA). Results of one-way ANOVA and post hoc testing by Dunnett’s multiple comparisons test are shown; * p-value ≤0.05. E. Representative images of in-situ hybridization using *c-fos* (white) and sytox orange to show nuclei (red) in the dorsomedial hypothalamus (DMH), indicated by white outline, for each group defined in A. Scalebar 500 µm. F. Quantification of the percentage of positive cells in the dorsomedial hypothalamus (DMH). Results of one-way ANOVA and post hoc testing by Dunnett’s multiple comparisons test are shown; ** p-value ≤0.005.

Surveying the thermoregulatory circuits and comparing *c-fos* expression over these timepoints revealed that the POA (Figure 2B) showed an approximately 0.5% increase in the proportion of *c-fos* positive cells after 40 hours of torpor compared to IBE-12h (Figure 2C & D, p < 0.01, one-way ANOVA, Dunnett’s multiple comparison test). Contrastingly, we saw significant changes in the number of *c-fos* positive cells in the DMH during the entry phase of the T-A cycle (2.25% positive cells, Figure 2E & F, p < 0.005, one-way ANOVA, Dunnett’s multiple comparison test). In the DMH during arousal *c-fos* positive cells were detected but these changes were variable and therefore not significant. We also surveyed other neuronal centres in the hypothalamus (Figure 2B; ARC and VMH), less than 1% of cells showed *c-fos* expression during torpor and almost no cells were positive in the transition phases of the T-A cycle (Supplementary Figure 2B & C).

### *c-fos* induction in choroid plexus, tanycytes and pars tuberalis and during early arousal

We noted high expression of *c-fos* in the choroid plexus (CP) (Figure 3A, B & C), with an average of 65% of CP cells showing positive staining in torpid animals, rising to 93% in the earliest arousing animals (AΔ0.5`°`C), compared to 0.3% of positive cells in IBE-12h (Figure 3C, IBE-12h versus AΔ0.5`°`C p=0.0005, IBE-12h versus T-40h p=0.0063, one-way ANOVA, post-hoc testing with Dunnett’s multiple comparisons). The circumventricular organs, the vascular organ of lamina terminalis (OVLT) and median eminence (ME) showed approximately 5% of cells as positive during torpor and arousal but there was high inter-individual variation within the groups (Supplementary Figure 3A-D). Whilst surveying the ME, we noted that the pars tuberalis (PT) of the adjacent pituitary stalk (Figure 3A) showed very high *c-fos* expression (22.6% of cells) at AΔ0.5`°`C (Figure 3D & E, IBE12h versus AΔ0.5`°`C p <0.0001, one-way ANOVA, post-hoc testing with Dunnett’s multiple comparisons). Similarly, alpha tanycytic cells lining the 3^rd^ ventricle of the mediobasal hypothalamus (αTC, Figure 3A), showed increased *c-fos* expression in AΔ0.5`°`C (7.3%, Figure 3F & G, IBE-12h versus AΔ0.5`°`C p=0.0064, one-way ANOVA, post-hoc testing with Dunnett’s multiple comparisons). Beta tanycytes lining the 3^rd^ ventricle but in closer proximity to the ME also showed an increase in *c-fos* positive cells but to a lesser degree (Supplementary Figure 3E & F). Tanycytes have long projections into the hypothalamus and down to the pars tuberalis (Guerra et al., 2010; Güldner and Wolff, 1973; Rodríguez et al., 2005), we used vimentin to stain these projections, showing infiltration to the DMH, VMH, ARC and PT (Figure 3H & I).

**Figure 3:**
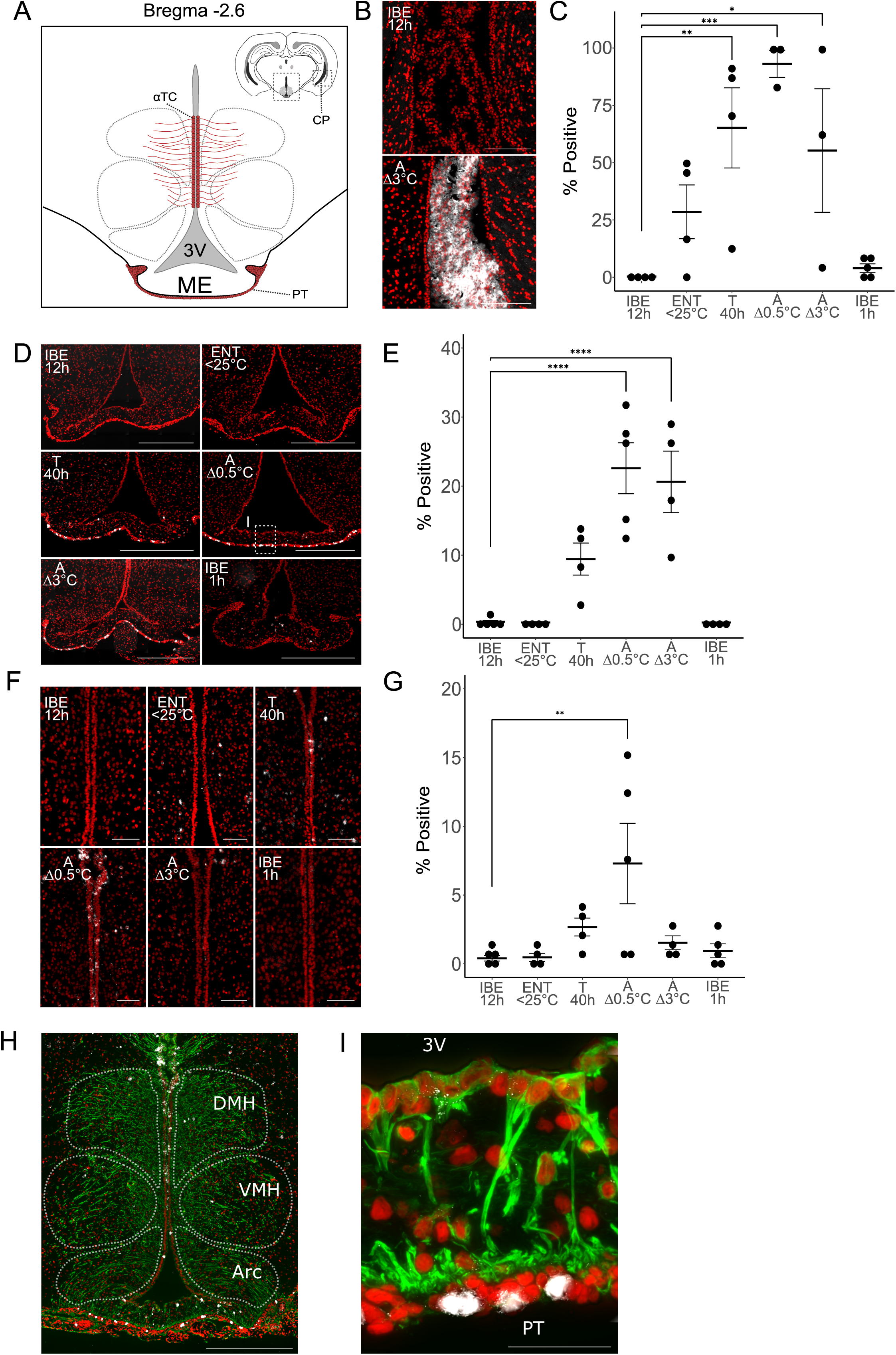
*c-fos* expression in the choroid plexus, pars tuberalis and tanycytes during early arousal from torpor. A. A schematic indicating Bregma −2.6 and the location of α-tanycytes (αTC), Pars tuberalis (PT) and choroid plexus (CP). ME; median eminence. Grey dotted lines indicate the regions of the hypothalamic nuclei. Note the CP is in the lateral ventricles and is therefore displayed in the inset relative to the hypothalamic region shown. B. Representative images of in-situ hybridization using *c-fos* (white) and sytox orange to show nuclei (red) in the choroid plexus (CP), for IBE-12h and AΔ3°C. Scalebar 500 µm. C. Quantification of the percentage of positive cells in the choroid plexus (CP). Results of one-way ANOVA and post hoc testing by Dunnett’s multiple comparisons test are shown; * p-value ≤0.05, ** p-value ≤0.005, *** p-value ≤0.0005 D. Representative images of in-situ hybridization using *c-fos* (white) and sytox orange to show nuclei (red) in the pars tuberalis (PT), for each group defined in Figure 2A. The representative image for AΔ0.5°C contains a white dotted line box indicating the zoomed in image shown in I. Scalebar 500 µm. E. Quantification of the percentage of positive cells in the pars tuberalis (PT). Results of one-way ANOVA and post hoc testing by Dunnett’s multiple comparisons test are shown; **** p-value <0.0001 F. Representative images of in-situ hybridization using *c-fos* (white) and sytox orange to show nuclei (red) in the α-tanycytes (αTC) region, for each group defined in Figure 2A. Scalebar 200 µm. G. Quantification of the percentage of positive cells in the α-tanycytes (αTC) region. Results of one-way ANOVA and post hoc testing by Dunnett’s multiple comparisons test are shown; ** p-value ≤0.005. H. Vimentin staining (green) for tanycyte process in the mediobasal hypothalamus. *c-fos* (white) and sytox orange to show nuclei (red). The hypothalamic nuclei are indicated by white dotted lines. DMH; dorsomedial hypothalamus, VMH; ventromedial hypothalamus, Arc; Arcuate nucleus. Scalebar 500 µm. I. A zoomed in image of the pars tuberalis (PT) shown in D during early arousal (AΔ0.5°C). Vimentin staining (green) for tanycyte processes, *c-fos* shown in white, and sytox orange to show nuclei is red. 3V; third ventricle. Scalebar 50 µm.

## Discussion

We have monitored spontaneous T-A cycles in the golden hamster at high temporal and temperature resolution to capture the initiation of spontaneous arousal from torpor. In two hypothalamic neuronal centres linked to thermoregulation, the POA and the DMH (Cannon and Nedergaard, 2004; Morrison, 2016), no significant change in *c-fos* expression was seen during spontaneous arousal, and the only significant change observed was increased expression during torpor entry in the DMH, potentially reflecting the tightly controlled cooling process (Florant and Heller, 1977). This contrasts with the pattern seen in 3 non-neuronal tissues, the CP, PT and tanycytes, in which we detected the highest level of *c-fos* RNA expression when BAT temperature had risen only 0.5°C above torpor baseline. This indicates that these regions are sensitive to physiological changes at an early stage in the arousal process. This extends similar findings in the thirteen lined ground squirrel (Bratincsák et al., 2007) and is consistent with involvement of these sites in metabolic feedback regulation of T-A cycling.

The expression of *c-fos* in non-neuronal cells in the choroid plexus, the mediobasal hypothalamus and the PT may reflect ionic changes that are not so clearly linked to “activation” as is the case for neuronal *c-fos* expression (Lara Aparicio et al., 2022), but are nonetheless interesting given the highly time- and tissue-restricted nature of this response. The primary function of the choroid plexus is to produce and control the composition of CSF, maintaining a blood-CSF barrier that gates the exchange of metabolites and factors into the brain (Lun et al., 2015). In the mouse, the CP shows pronounced daily changes in its production of major transport proteins, notably the thyroid hormone carrier protein transthyretin (Duarte et al., 2020; Fame et al., 2023), while accumulation of hibernation factor in the CP of Siberian chipmunks shows clear circannual variation (Kondo et al., 2006). These observations suggest that CP function changes over time, correlated with daily and seasonal changes in brain energy demand.

This framing of CP function has clear parallels with the function of tanycytes in the mediobasal hypothalamus: These modified glial cells line the walls of the 3^rd^ ventricle, and serve a gating function modulating the influence of nutrients, hormones and metabolites as feedback signals on neuronal populations involved in appetite, energy balance and reproduction (Balland et al., 2014; Bolborea et al., 2020; Frayling et al., 2011; Langlet, 2014; Langlet et al., 2013; Lhomme et al., 2021). Tanycytes have been linked to circadian control of glucose homeostasis (Rodríguez-Cortés et al., 2022) and have emerged as key cellular substrate for seasonal modulation of hypothalamic function (Dardente et al., 2014; Dardente et al., 2019; Hazlerigg and Loudon, 2008). This latter aspect depends on photoperiod-dependent production of thyrotropin by neighbouring PT cells which in turn acts via tanycytic TSH-receptor expression (Hanon et al., 2008). This then modulates tanycytic uptake of thyroid hormone by monocarboxylate transporter 8 (SLC16A2) (Friesema et al., 2003; Petri et al., 2016) and tanycytic conversion of thyroid hormone by deiodinases (dio2 / dio3) (Dardente et al., 2014; Dardente et al., 2019; Hazlerigg and Loudon, 2008). The relevance of this pathway for expression of torpor is suggested by the blockade of daily torpor expression in Siberian hamsters by exogenous thyroid hormone delivery to the DMH (Murphy et al., 2012), a site to which tanycyte cell processes project (Figure 3H) (Guerra et al., 2010; Güldner and Wolff, 1973; Rodríguez et al., 2005). Furthermore, in the Arctic ground squirrel, circannual termination of hibernation correlates with increased deiodinase 2 gene expression in the mediobasal hypothalamus (Chmura et al., 2022).

Hence it can be seen that involvement in temporal changes in metabolic energy demand and thyroid metabolism constitutes a unifying functional framework for the 3 non-neuronal tissues in which we see peak *c-fos* expression at arousal initiation. This leads us to hypothesise that *c-fos* induction occurs in response to metabolic changes developing during torpor, reflecting the direct sensitivity of these tissues to changing concentrations of blood- or CSF-borne metabolites. Further experiments are required to test this hypothesis and to explore how changes in tanycyte or CP activity might couple to the neural circuits driving thermogenesis at arousal.

## Acknowledgements

The authors would like to thank our animal technicians Hans Lian, Hans-Arne Solvang and Renate Thorvaldsen. The authors also thank Valerie Simonneaux and Paul Klosen for the gift of the *c-fos* plasmid. We acknowledge the UiT core microscopy facility for the use of the confocal microscope.

## Data availability

The analyzed data and code for figures for this study can be found in GitHub repository https://github.com/ShonaWood/cFOSGHam

---

**Supplementary Figure 1: Physiological monitoring of hibernation in the golden hamster**

A. Percentage of animals hibernating in response to SP-cold conditions over time.

B. Core body temperature (T_b_) in response to the transition from long photoperiod (LP) at ambient temperature (T_a_) 21`°`C, to short photoperiod (SP) at Ta 21`°`C (first dotted horizontal line), to SP and T_a_ 6.5`°`C. Prior to T-A cycling diel patterning of mean core body temperature was observed at all stages of the experiment, except the first 4 weeks of cold where the T_b_ is reducing (see inset periodograms). Periodograms were generated using ActogramJ and the chi-square method.

C. Raleigh plots showing the time of test-drop entry and arousal from 24 individuals.

Photoperiod is represented by grey for dark and yellow for light. The black dots are an individual arousal or entry event. The black arrow indicates the mean time of entry or arousal, and the light grey arrows indicates the standard deviation. The length of the arrow represents mean resultant length as an indicator of concentration around the mean, therefore reflecting the statistical significance. The Raleigh statistical was used to test for a time-of-day preference in arousal and entry. Both entry and arousal were significant p<0.05.

D. Top: Duration (hours) of inter bout euthermia after previous corresponding torpor bout for 22 individuals; listed on the side with individual colours shown in each box plot. Bottom: Torpor bout duration (hours) for each torpor bout made.

**Supplementary Figure 2: Validation of *c-fos* probe and numbers of *c-fos* positive cells in the ARC and VMH**

A. In-situ hybridization of the suprachiasmatic nucleus (SCN) in the golden hamster for *c-fos* RNA (brown staining). Left: no light pulse during the dark phase. Right: 20-minute light pulse during the dark phase. 3V; 3^rd^ ventricle, indicated by black dotted line.

B. Quantification of the percentage of positive cells in the arcuate nucleus (ARC) region. Results of one-way ANOVA and post hoc testing by Dunnett’s multiple comparisons test are shown; * p-value ≤0.05.

C. Quantification of the percentage of positive cells in the ventromedial hypothalamus (VMH) region. Results of one-way ANOVA and post hoc testing by Dunnett’s multiple comparisons test are shown; ** p-value ≤0.005.

**Supplementary Figure 3: *c-fos* expression in the OVLT, ME and beta tanycytes**

A. Quantification of the percentage of positive cells in the Vascular organ of lamina terminalis (OVLT) region. Results of one-way ANOVA and post hoc testing by Dunnett’s multiple comparisons test are shown; ** p-value ≤0.005.

B. Representative images of in-situ hybridization using *c-fos* (white) and sytox orange to show nuclei (red) in the OVLT region, for IBE-12h and AΔ3°C. Vimentin staining is shown in green. Scalebar 200 µm.

C. Quantification of the percentage of positive cells in the median eminence (ME). Results of one-way ANOVA and post hoc testing by Dunnett’s multiple comparisons test are shown; ns = not significant.

D. Representative images of in-situ hybridization using *c-fos* (white) and sytox orange to show nuclei (red) in the ME region, for IBE-12h and AΔ3°C. Vimentin staining is shown in green. Scalebar 100 µm.

E. Quantification of the percentage of positive cells in the β-tanycytes (βTC) region. Results of one-way ANOVA and post hoc testing by Dunnett’s multiple comparisons test are shown; * p-value ≤0.05.

F. Representative images of in-situ hybridization using *c-fos* (white) and sytox orange to show nuclei (red) in the β-tanycytes (βTC) region, for IBE-12h and AΔ3°C. Vimentin staining is shown in green. Scalebar 500 µm.

## Author Contributions

Conceptualization: S.H.W., D.G.H., F.A.F.M., Methodology: F.A.F.M., F.C.M., V.J.M, Software: F.A.F.M., Validation: F.A.F.M., F.C.M., Formal analysis: F.A.F.M., S.H.W., D.G.H., Investigation: F.A.F.M, F.C.M., V.J.M., Resources: S.H.W., Data curation: F.A.F.M., Writing - original draft: F.A.F.M., S.H.W., D.G.H., Writing - Review & editing: All authors, Visualization: F.A.F.M., Supervision: S.H.W., D.G.H., Project administration: S.H.W., Funding acquisition: S.H.W.

