## Supplementary figures and images for "*c-fos* induction in the choroid plexus, tanycytes and pars tuberalis is an early indicator of spontaneous arousal from torpor in a deep hibernator"

### Supplementary figure 1

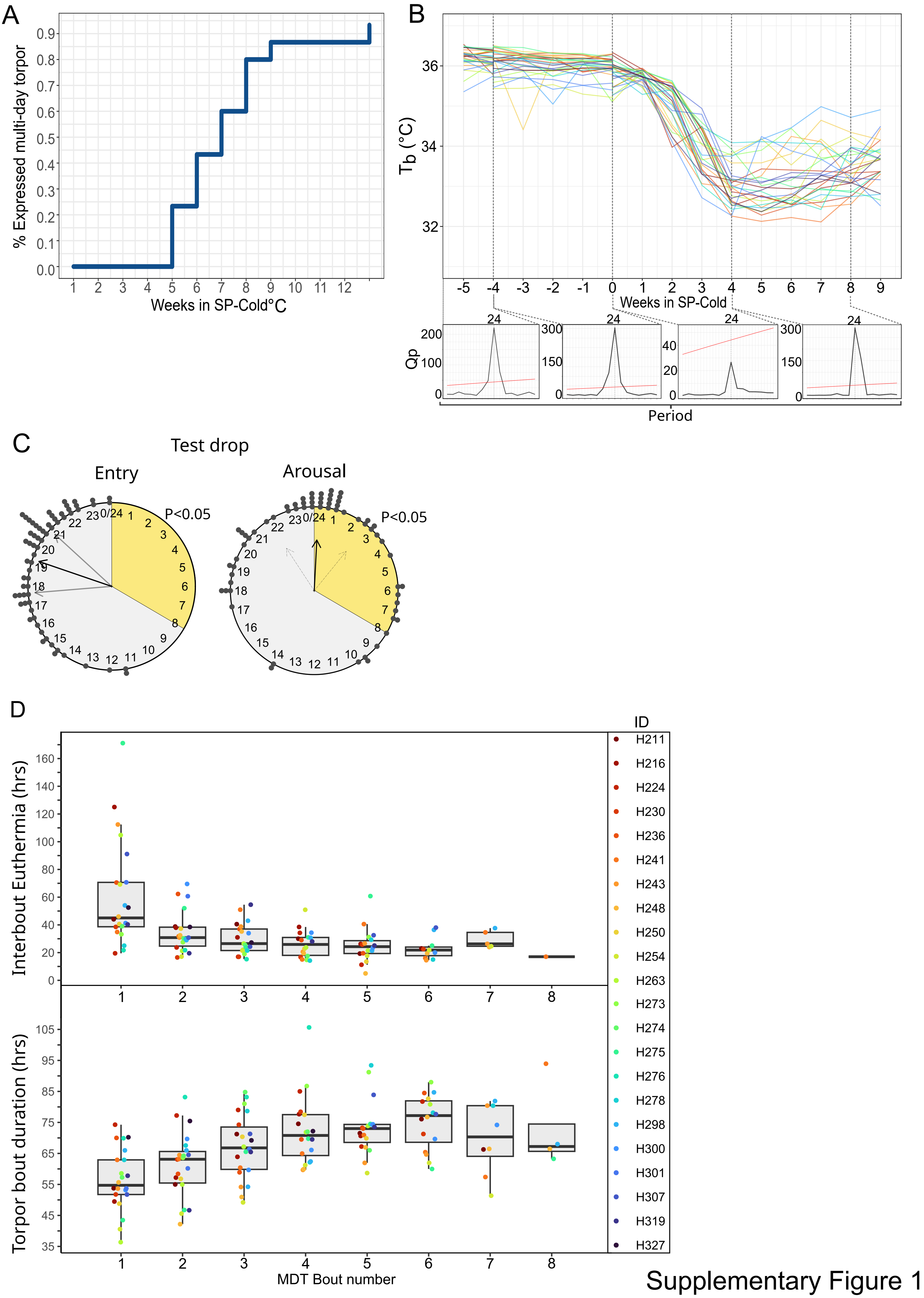


Supplementary Figure 1

### Supplementary figure 2

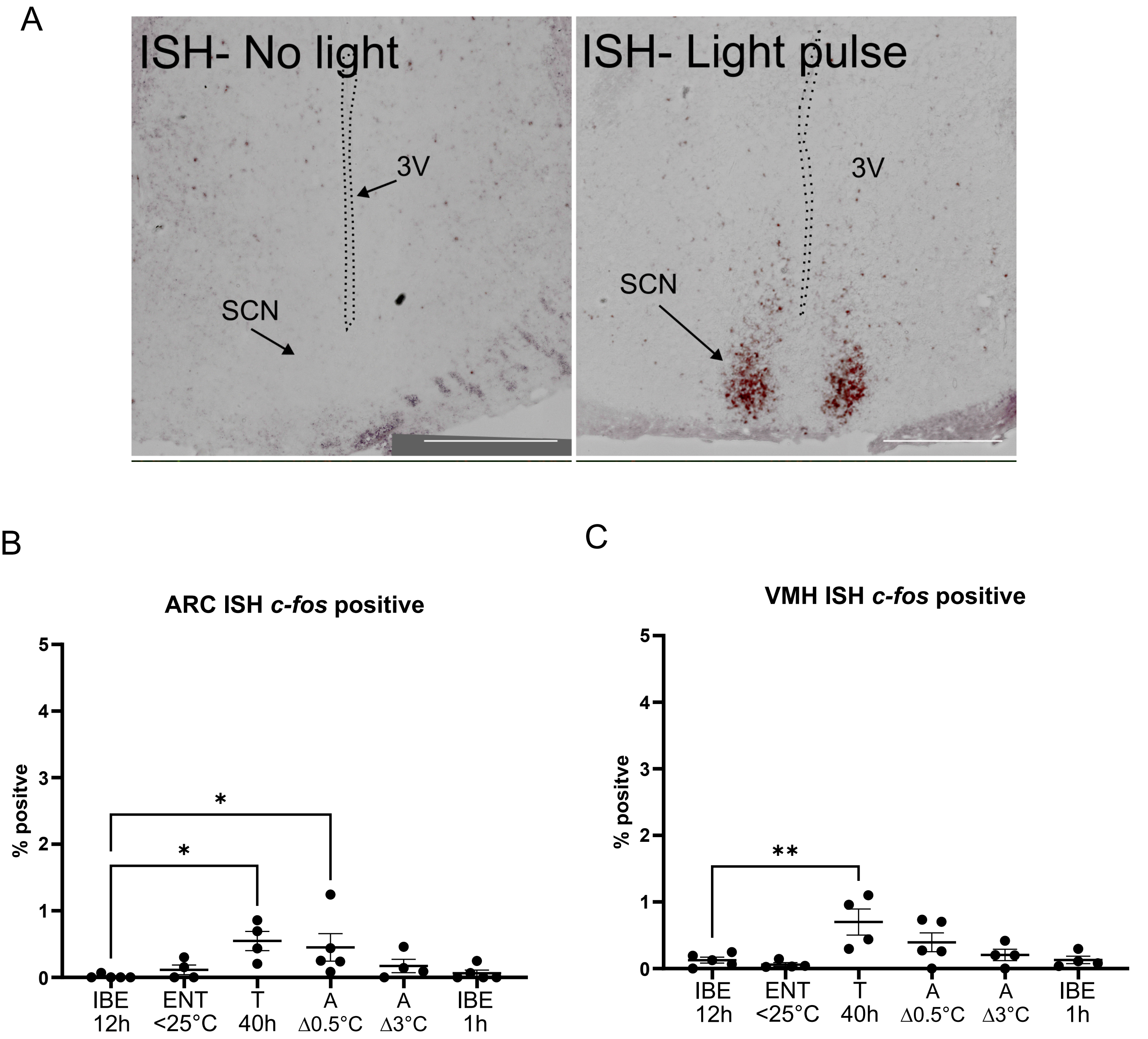


Supplementary Figure 2

### Supplementary figure 3

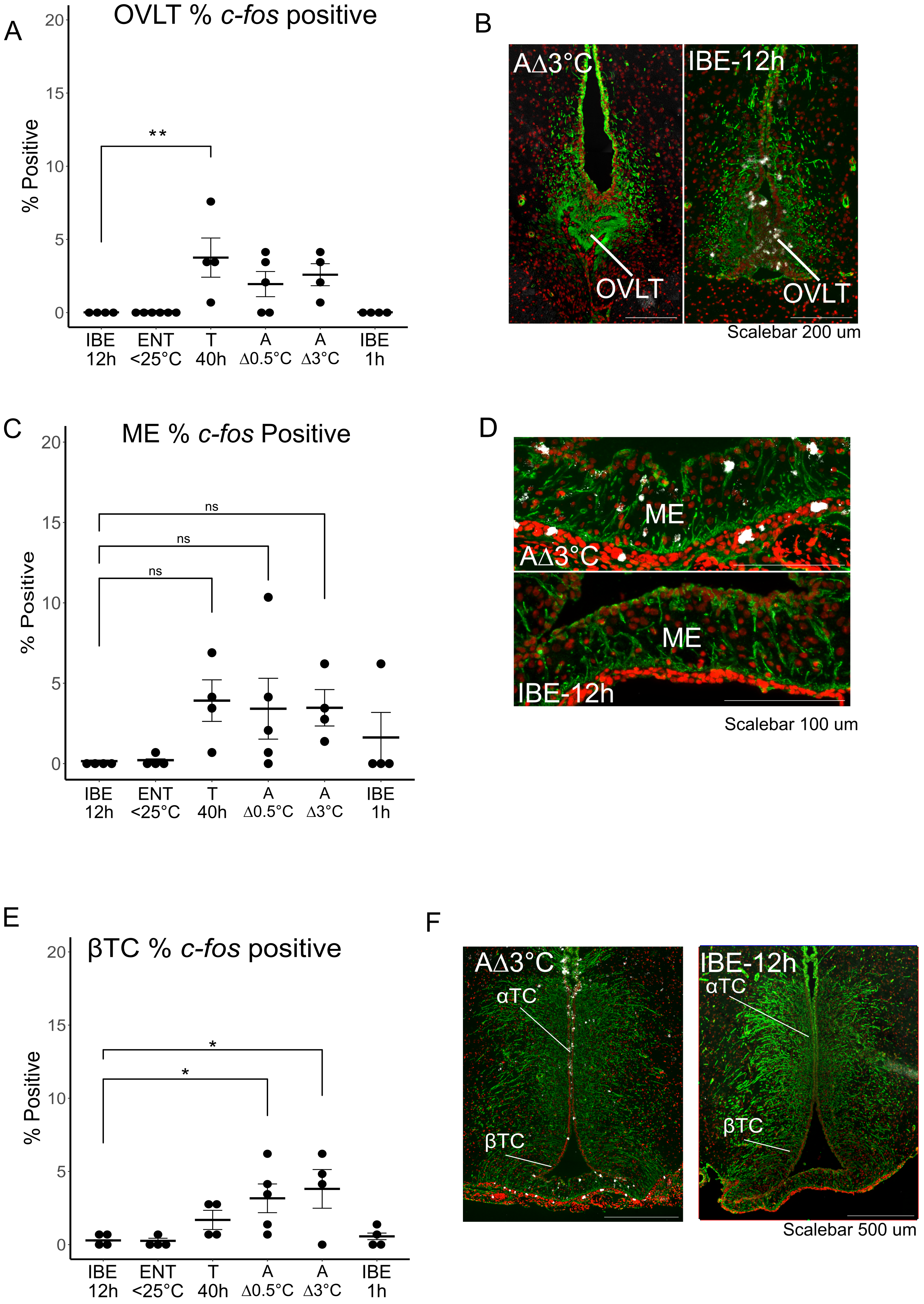


Supplementary Figure 3
